# Protected area planning to conserve biodiversity in an uncertain future

**DOI:** 10.1101/2022.11.18.517054

**Authors:** Richard Schuster, Rachel Buxton, Jeffrey O. Hanson, Allison D. Binley, Jeremy Pittman, Vivitskaia Tulloch, Frank A. La Sorte, Patrick R. Roehrdanz, Peter H. Verburg, Amanda D. Rodewald, Scott Wilson, Hugh P. Possingham, Joseph R. Bennett

## Abstract

Protected areas are a key instrument for conservation. Despite this, they are vulnerable to risks associated with weak governance, land use intensification, and climate change. Using a novel hierarchical optimization approach, we identified priority areas for expanding the global protected area system to explicitly account for such risks whilst maximizing protection of all known terrestrial vertebrate species. We illustrate how reducing exposure to these risks requires expanding the area of the global protected area system by 1.6% while still meeting conservation targets. Incorporating risks from weak governance drove the greatest changes in spatial priorities for protection, while incorporating risks from climate change required the largest increase in global protected area. Conserving wide-ranging species required countries with relatively strong governance to protect more land when bordering nations with comparatively weak governance. Our results underscore the need for cross-jurisdictional coordination and demonstrate how risk can be efficiently incorporated into conservation planning.

**Article Impact Statement:** Accounting for governance, land use and climate risks will result in more resilient and effective conservation effort for biodiversity.

## Introduction

Protecting land is one of the best strategies to stem the biodiversity crisis (Watson et al. 2014) and a cornerstone of international agreements to safeguard biodiversity (CBD 2010, 2020). The effectiveness of a global network of protected areas depends upon the identification of areas that will both meet the needs of species and provide the greatest return on conservation investments. Yet most spatial planning efforts base decisions heavily upon the estimated ecological value of land (Brooks et al. 2006; CBD 2010, 2020; Venter et al. 2014) and carry the tacit, but often incorrect, assumption that protection will be enforced, effective, and permanent. However, there is strong evidence that protected areas are subject to risks associated with weak governance (e.g., political instability and corruption; Schulze et al. 2018), land use intensification (e.g., deforestation and degazetting of parks; Tesfaw et al. 2018), and climate change (e.g., extreme weather events; Maxwell et al. 2019). There is also evidence that patterns in protected area vulnerability vary spatially, with some regions and jurisdictions being more vulnerable to protected area degazettement or degradation than others (Mascia & Pailler 2011; Leberger et al. 2020). Few conservation plans have explicitly considered these risks (McBride et al. 2007; Alagador et al. 2014). Here we demonstrate how we can expand the planning lens to include risks as well as ecological value, with the goal of improving the resilience and performance of protected areas.

We considered the following three broad categories of risk, which we defined as factors likely to diminish the long-term effectiveness of protected areas: (i) governance, (ii) land use, and (iii) climate. We then generated plans for establishing protected areas (“prioritizations”) based on scenarios for different risk factors. Our framework provides a flexible approach for incorporating multiple risk metrics into conservation decision making.

## Methods

We considered the influence of risk categories on allocating protection decisions at a global scale in suitable habitat for all 29,350 vertebrate species from the IUCN Red List of Threatened Species (IUCN 2019) using a multi-objective optimization approach. To incorporate risk categories, we built on the minimum set problem, where the objective is to reach species distribution protection targets while accounting for one constraint such as land cost or area (Margules & Pressey 2000; Moilanen et al. 2009). We expanded this approach to include multiple objectives accounting for risk in the problem formulation, by treating each risk layer as a separate objective in the problem formulation (Deb 2014).

### Biodiversity Data

We produced area of habitat (AOH) estimates for 10,774 species of birds, 5,219 mammals, 4,462 reptiles and 6,254 amphibians with available IUCN range polygon data following the procedure outlined in Brooks et al. (2019). Species range polygons obtained from the IUCN Red List spatial data (https://www.iucnredlist.org/) and Birdlife International (http://datazone.birdlife.org/species/requestdis) were first filtered for ‘extant’ range then rasterized to a global 1 km grid in the Eckert IV equal area projection. Individual species range rasters were then modified to only include land cover classes that match the habitat associations for each species. Habitat associations were obtained from the IUCN Red List species habitat classification scheme and were matched to ESA land cover classes for the year 2018 following Santini et al. (2019). Habitat classifications of both ‘suitable’ and ‘marginal’ were used and included those identified as major importance. ESA land cover classification data was aggregated from 300 m resolution to match the global 1 km grid using a majority rule. Species ranges were additionally filtered so that only areas within a species’ accepted elevational range were included. Elevation limits were obtained from IUCN Red List entries for each species. Global elevation data derived from SRTM was obtained from WorldClim v. 2 (Fick & Hijmans 2017). For bird species with multiple seasonal distributions, data for resident, breeding, and non-breeding ranges were processed separately. For each AOH dataset, we then calculated the proportion of suitable habitat at a 10 x 10 km resolution which was the resolution used in the optimization analyses.

### Basic administrative delineations

National boundaries were derived from the Global Administrative Areas database (http://gadm.org). We obtained protected area boundaries from the World Database on Protected Areas (https://www.protectedplanet.net; (UNEP-WCMC & IUCN 2020)). Following standard procedures for cleaning the protected area dataset (Butchart et al. 2015), we (i) projected the data to an equal-area coordinate system (World Behrman), (ii) excluded reserves with unknown or proposed designations, (iii) excluded UNESCO Biosphere Reserves (Coetzer et al. 2014), (iv) buffered sites represented as point localities according to their reported area in the database (see UNEP-WCMC & IUCN (2020) for buffer sizes), (v) dissolved boundaries to prevent issues with overlapping areas, and (vi) removed slivers (code available at https://github.com/jeffreyhanson/global-protected-areas). After the protected area data were modified as described above, we overlaid the protected area boundaries with a 10 x 10 km grid covering the Earth and coded grid cells as protected if the protected area covered >50% of the cell following common practice (e.g. Hanson et al. 2020). These spatial data procedures were implemented using ArcMap (version 10.3.1) and python (version 2.7.8).

### Governance risk

We used worldwide governance indicators from the World Bank to capture governance risk (Kaufmann et al. 2011). The indicators include six scaled measures: voice and accountability; political stability and absence of violence; government effectiveness; regulatory quality; rule of law; and control of corruption (see Table S1 for definitions). We chose these indicators because evidence suggests that they reliably predict protected area effectiveness (Barnes et al. 2016) and state investment and efforts for biodiversity conservation (Coetzer et al. 2014). For each country, we used a mean of annual averages of all six measures (Baynham-Herd et al. 2018) (Figure S1).

### Land use risk

We used a global land systems map produced by Kehoe et al. (2017) to incorporate the risk of land-use change. This map is based on a global land systems map for the year 2000 (Asselen & Verburg 2012) at a 9.25 km^2^ spatial resolution but is refined based on recent land-cover and land-use datasets to a spatial resolution of 1 km^2^. Kehoe et al. (2017) further estimated the impact of land use and land use intensity on biodiversity, with data originating from the PREDICTS project (Hudson et al. 2014). They first matched their land-systems classes to varying intensity levels for each land use type (for detailed conversion table, see Asselen & Verburg 2012). This allowed Kehoe et al. (2017) to calculate average biodiversity loss per land system (relative to an unimpacted standard) by taking the mean model estimates of biodiversity loss per land use intensity class from previous work (Hudson et al. 2014). The result gives average relative biodiversity gain or loss per land-system class. Here, we used their modelled mean estimates (following Newbold et al. (2015)) of relative percent biodiversity change for each land-system class for species abundance as a measure of the land use pressure (Figure S2).

### Climate risk

We used velocity of climate change, which is an instantaneous measurement of how projected temperature increases translate to horizontal climate velocity on the landscape (Loarie et al. 2009). It is an integration of both the rate of change in average climate and landscape properties that govern how bands of similar temperature redistribute spatially as climate changes. For example, in a region with high topographic diversity, a species may be able to track its climatic niche through relatively small dispersal distances (e.g. 10s or 100s of meters) upslope or downslope. By contrast, keeping pace with preferred climate under the same magnitude of temperature rise in the plains may require much larger dispersal distances – 100s or 1000s of kilometers. Velocity of future temperature change used here follows the method of Loarie et al. (2009), and is essentially the ratio of the projected temporal rate of change (C/year) to the spatial rate of change (C/km). Projected temporal rate of change was based on the 20-year mean (2040-2060) projection for mean annual temperature from the HadGEM2-ES model (CMIP5) and the baseline (1960-1990) temperature available from Worldclim v1.4. Spatial rate of change was derived from 30 arc second elevation data and calculated with the ‘terrain’ function from the R ‘raster’ package. We also explored an alternative measure of climate risk: exposure to extreme events. Detailed methods and results for this alternative measure are provided in the online Supporting Information.

### Multi-objective optimization of risk reduction

We created 15 planning scenarios, such that solutions accounted for all possible combinations of risk categories within each hierarchical level (Table 1). We then compared these risk-based solutions to those produced with a baseline scenario that adopted the traditional area-minimizing approach to optimization without considering risk.

**Table 1.** Scenarios explored and global protection results. The risk factor order represents the order risk factors were included in the hierarchical prioritization. (G = governance, L = land use, C = Climate).

| Scenario | Risk factors included | Global land area protected [%] |
| --- | --- | --- |
| null | - | 21.27 |
| 1 | G | 21.35 |
| 2 | L | 22.31 |
| 3 | C | 23.79 |
| 4 | G > L | 21.93 |
| 5 | L > G | 22.18 |
| 6 | G > C | 23.78 |
| 7 | C > G | 23.31 |
| 8 | L > C | 23.52 |
| 9 | C > L | 22.99 |
| 10 | G > L > C | 23.52 |
| 11 | G > C > L | 23 |
| 12 | L > G > C | 23.5 |
| 13 | L > C > G | 23.08 |
| 14 | C > G > L | 22.3 |
| 15 | C > L > G | 22.99 |

We processed all data described previously to a 10 x 10 km resolution and clipped data to the extent of land based on the global administrative areas database to use as planning units in the optimization analyses. For biodiversity data, we calculated the proportion of suitable habitat in each 10 x 10 km pixel, for governance risk and land use risk we used the nearest neighbor approach, and for climate risk we calculated the mean. We used this resolution as a tradeoff between precision and computational feasibility. Our multi-objective approach uses a hierarchical (lexicographic) framework that assigns a priority to each objective, and sequentially optimizes for the objectives in order of decreasing priority. At each step, it finds the best solution for the current objective, but only from among those that would not degrade the solution quality for higher-priority objectives. We considered up to three objectives in our prioritization scenarios, i) governance risk, ii) land use risk, and iii) climate risk. To compare different scenarios, we calculated solutions for each unique objective combination (n = 15), as well as one where we use a constant objective function as the baseline scenario, as the order of the hierarchy can influence the results (See Table 1 for details).

In systematic conservation planning, conservation features describe the biodiversity units (e.g., species, communities, habitat types) that are used to inform protected area establishment. Planning units describe the candidate areas for protected area establishment (e.g., cadastral units). Each planning unit contains an amount of each feature (e.g., presence/absence, number of individuals). A prioritization describes a candidate set of planning units selected for protected establishment. Each feature has a representation target indicating the minimum amount of each feature that ideally should be held in the prioritization (e.g., 50 presences, 200 individuals). To minimize risk, we have a set of datasets describing the relative risk associated with selecting each planning unit for protected area establishment. Thus, we wish to identify a prioritization that meets the representation targets for all of the conservation features, with minimal risk.

Let I denote the set of conservation features (indexed by <u>i</u>), and J denote the set of planning units (indexed by j). To describe existing conservation efforts, let p_j_ indicate (i.e., using zeros and ones) if each planning unit j ∈ J is already part of the global protected area system. To describe the spatial distribution of the features, let A_ij_ denote (i.e., using zeros and ones) if each feature is present or absent from each planning unit. To ensure the features are adequately represented by the solution, let t_i_ denote the conservation target for each feature i ∈ I. Next, let D denote the set of risk datasets (indexed by d). To describe the relative risk associated with each planning unit, let R_dj_ denote the risk for planning units j ∈ J according to risk datasets d ∈ D.

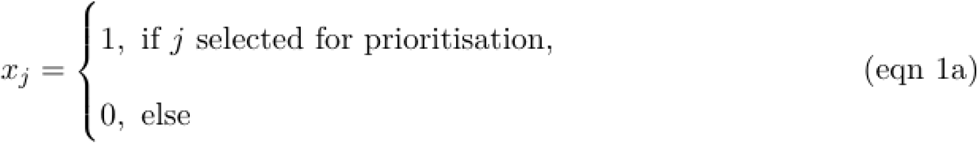

The problem contains the binary decision variables xj for planning units j ∈ J.

The reserve selection problem is formulated following:

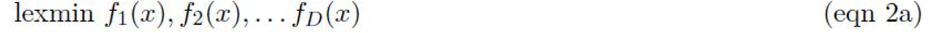

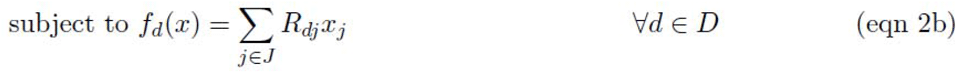

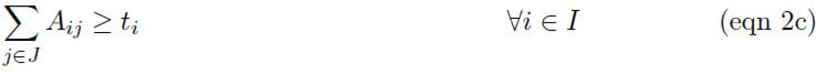

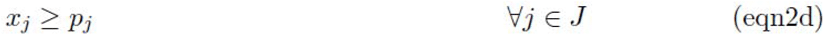

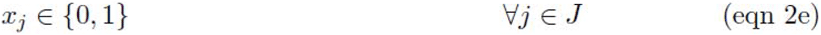

The objective function (eqn 2a) is to hierarchically (lexicographically) minimize multiple functions. Constraints (eqn 2b) define each of these functions as the total risk encompassed by selected planning units given each risk dataset. Constraints (eqn 2c) ensure that the representation targets (t_i_) are met for all features. Constraints (eqn 2d) ensure that the existing protected areas are selected in the solution. Finally, constraints (eqns 2e) ensure that the decision variables x_j_ contain zeros or ones.

For all scenarios we locked in current protected areas. Following Hanson et al. (2020), we used flexible targets for suitable habitat based on species’ ranges. Species with less than 1,000 km^2^ of suitable habitat were assigned a 100% target (1,802 amphibians, 893 avian, 645 mammalian, and 1,707 reptile species), species with more than 250,000 km2 of suitable habitat were assigned a 10% target (712 amphibians, 4,518 avian, 1,868 mammalian, and 595 reptile species) and species with an intermediate amount of suitable habitat were assigned a target by log-linearly interpolating values between the previous two thresholds (2,683 amphibians, 5,190 avian, 2,557 mammalian, and 2,160reptile species). Migratory bird species were assigned targets for each seasonal distribution separately. Additionally, to prevent species with very large suitable habitats from requiring excessively large amounts of area to be protected, the targets for species’ distributions larger than 10,000,000 km^2^ were capped at 1,000,000 km^2^. This upper limit affected only 206 (1%) species, and sensitivity analyses in a similar study showed that it had a negligible effect on results (Extended Data Fig. 1 in Hanson et al. 2020). We acknowledge that these targets are arbitrary; however, they are more precise than previous targets based on species’ ranges (which can contain a large amount of unsuitable habitat), and account for the increased vulnerability of species with smaller range sizes (Pimm & Raven 2000), as well as the difficulty in conserving all habitat for species that occur over large areas. We also acknowledge that we do not consider all sources of uncertainty, such as uncertainty in species distributions, or climate predictions. Rather, we focus on the risk categories identified above.

## Results

Scenarios that incorporated combinations of the three risk categories increased the priority area by only 1.6% on average (0.08 – 2.52%) compared to the baseline scenario based solely on ecological value to species. Among single-risk scenarios, accommodating risks due to climate change velocity required the greatest increase in global protected area, compared to scenarios including only governance or land use intensification risks (Table 1).

Scenarios shared many overlapping spatial priorities, which can be considered as reliably good investments in terms of both ecological value and risk management. Most notably, all 15 non-baseline scenarios prioritized the same 8.5 million km^2^ (5.8% of global land area) (“no regrets” areas, Fig. 2), much of which was located in western South America and southeast Asia (Figure S4). There was also substantial overlap among the priorities across scenarios within Conservation International’s global biodiversity hotspots (Myers et al. 2000), but many high overlap areas lie either outside of (53.3%) or in small areas within hotspots (Figure S5).

**Figure 1:**
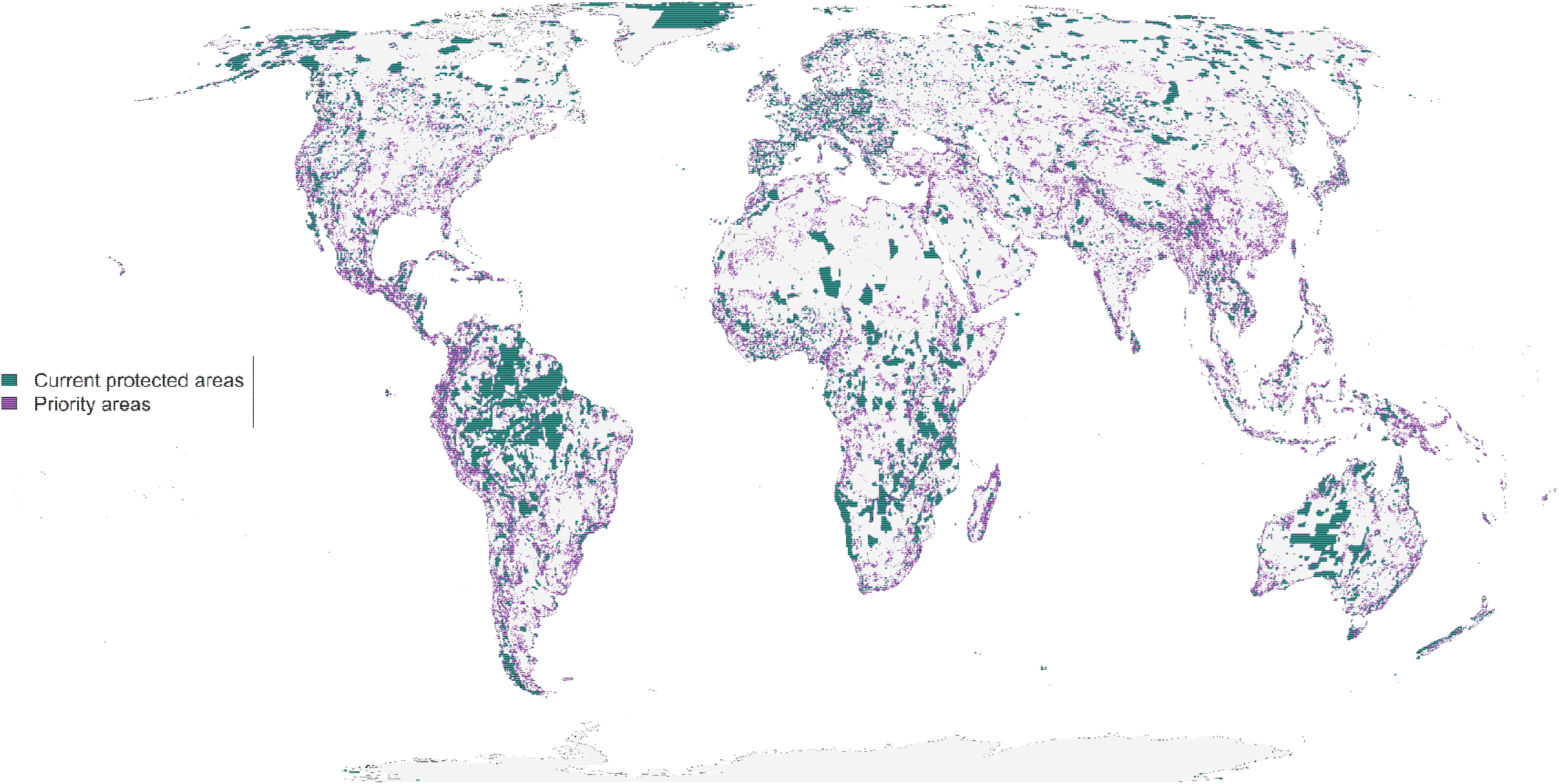
Spatial representation of priority areas for protection to account for governance, land use and climate risk together and in that order. Accounting for these risks to protected area effectiveness to produce more resilient conservation networks would require 23.5% of land surface to reach suitable habitat protection goals (*26*) for vertebrate species from the IUCN Red List of Threatened Species (*20*).

**Figure 2:**
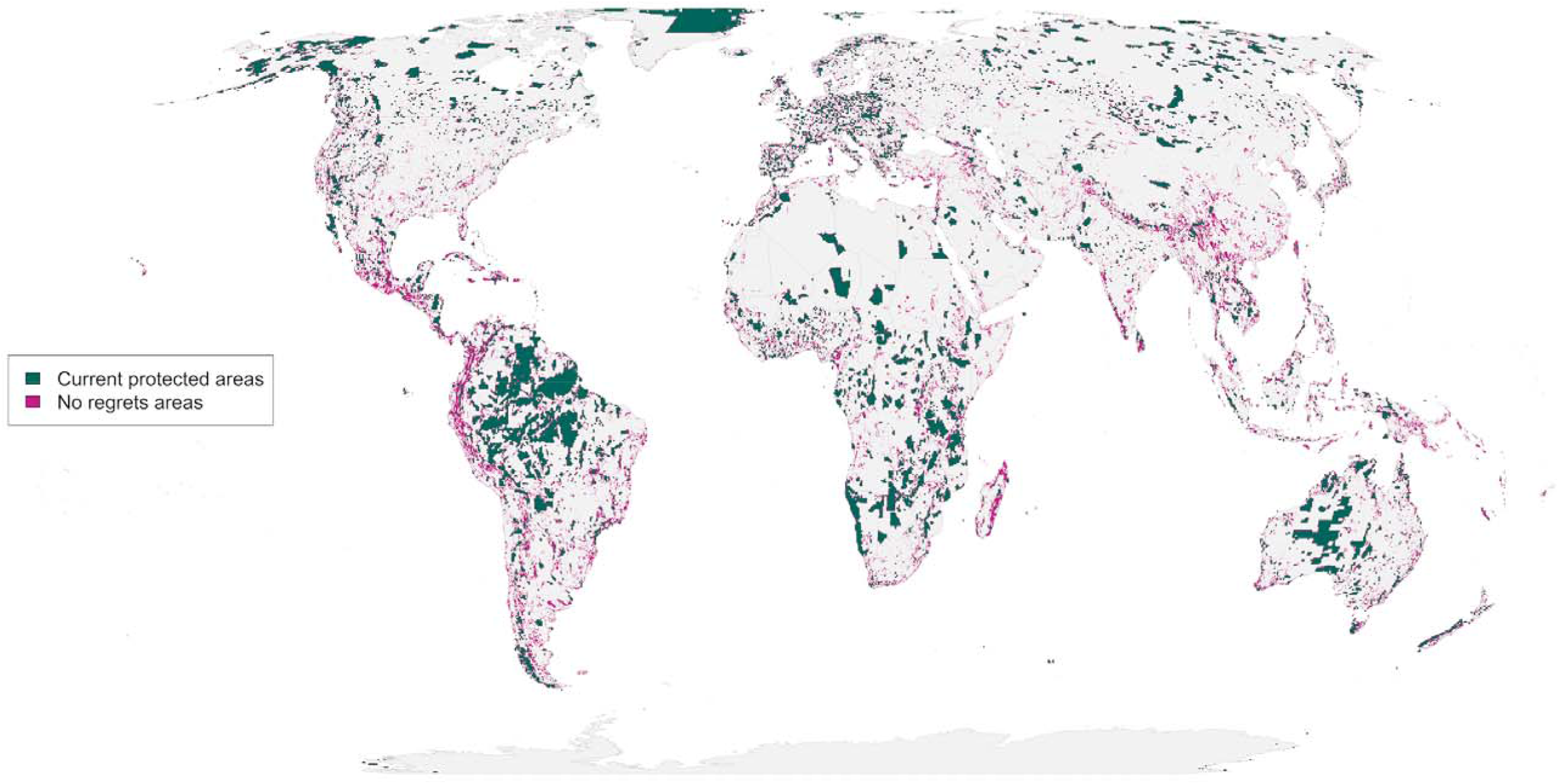
“No regrets” areas comprising 8.5 million km^2^ of land that was identified as priority habitat for protection regardless of the risks included in our analysis.

At the same time, risk scenarios elicited several prominent shifts in spatial priorities among areas varying in risk exposure (Fig. 3; Table S2). In some cases, high risks to protected areas in weakly-governed countries could be compensated by expanding protected areas in well-governed neighboring countries (Fig S6). For example, challenges to the transborder conservation of the wide-ranging and IUCN-vulnerable caribou (*Rangifer tarandus*) due to weak governance in Russia (Table S3) were mitigated by increasing the land area protected by Finland from 16.2% to 36.4% (Fig. 4). High exposure to risks from land use change could be offset in a similar fashion, such as by protecting more land in Liberia (32% versus 22.5% in the baseline scenario) than in the agriculturally intensifying nation of Sierra Leone (Fig. 4). Likewise, climate-associated risks in Hungary and Serbia (Figure S3) might be tempered by protecting twice as much land (20.4% versus 10.2% in baseline) in nearby Kosovo, which has lower predicted climate velocity (Fig. 4). Addressing risks from extreme weather events (La Sorte et al. 2021) (Figures S7 – S9) also required shifting some priority areas to less climatically volatile locations. Combining both climate velocity and extreme weather events into one metric illustrates a somewhat smoothed response (Figures S10 - S12).

**Figure 3:**
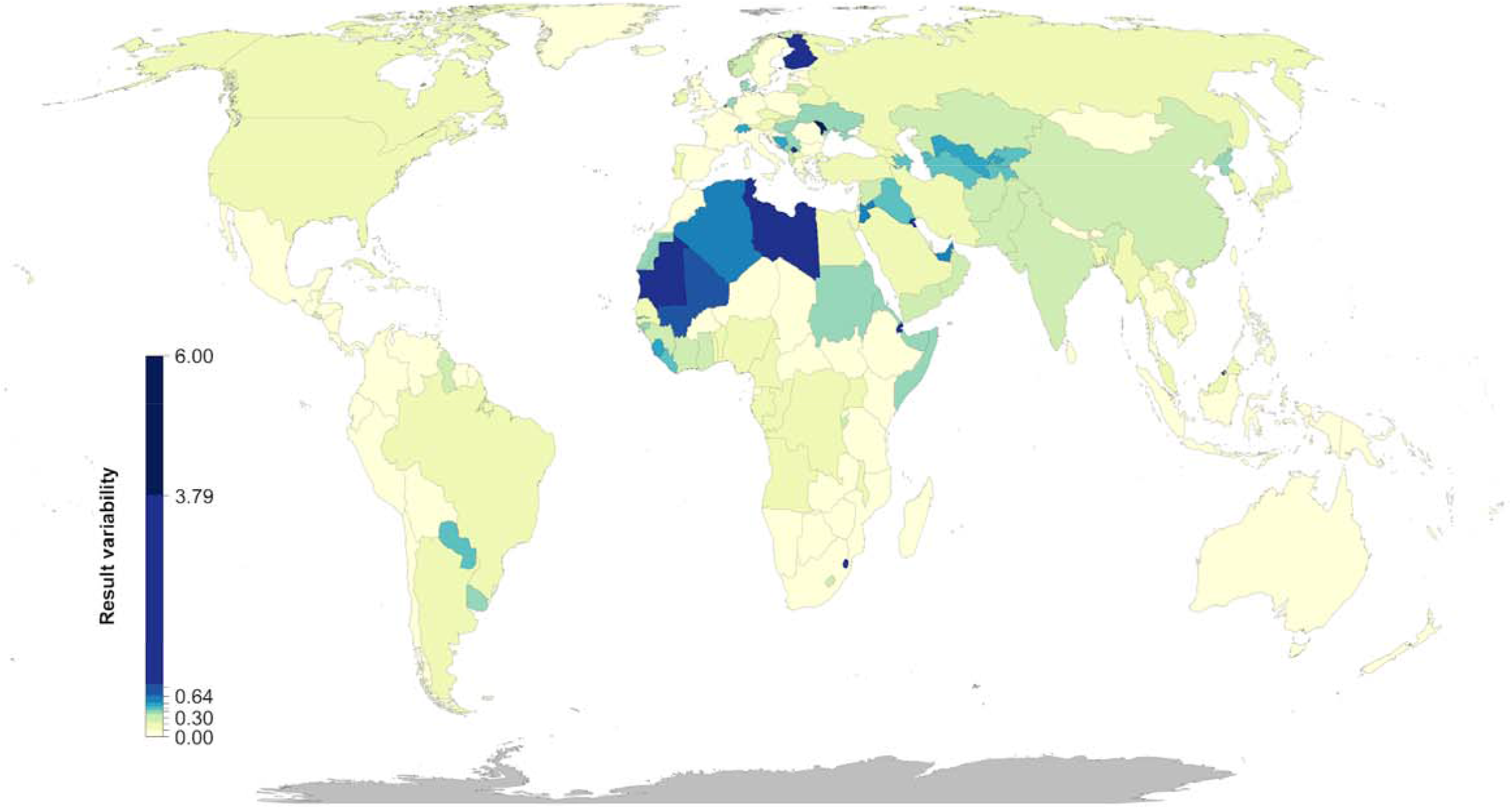
Percent country-level variation between the baseline scenario and the 15 scenarios including risk. Countries whose results are consistent across the 15 scenarios (e.g., Mexico) have low variation, while countries whose results are less consistent across the 15 scenarios (e.g., Finland) have high variation.xs

**Figure 4:**
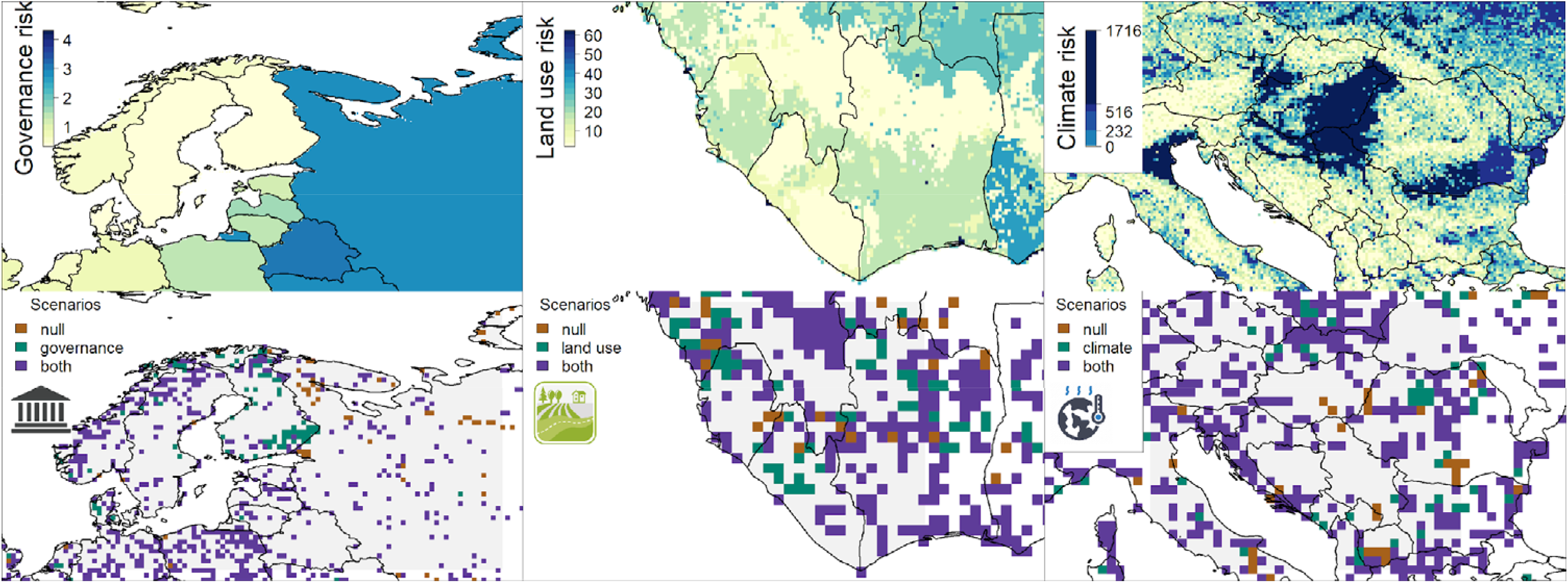
Contrast of using individual risk objectives (governance, land use, climate) to the baseline scenario of uniform objective structure. The top panels represent the individual risk data for the focal regions including northern Europe (left), west Africa (center) and southern Europe (right). In the bottom panels brown shows the baseline scenario, green the specific risk objective scenario results, and purple where both scenarios agree. The figures show how the spatial configuration of the solutions changes when risk is considered in a scenario. The top row of the maps represent data in their original resolution, the bottom row represents scenario results at a 10 x 10 km resolution.

## Discussion

Although a growing body of literature shows that protected areas are seldom effective if subject to unmitigated risks from land use change, weak governance and climate change (Schulze et al. 2018; Tesfaw et al. 2018; Maxwell et al. 2019), most systematic conservation planning efforts prioritize land based on ecological value and some measure of cost. We show how relatively small (1.6%) increases in land area, but importantly a change in the spatial configuration of where protected areas are placed, may reduce the vulnerability of protected areas to future threats if risk is explicitly considered during planning stages (Fig. 1). Across all planning scenarios, we identified 8.5 million km^2^ of priority lands that either uniquely contributed to conservation targets (e.g., high endemism) or were resilient to the risks we modeled. Countries with large proportions of land already in protection (e.g., Brazil with >30%) also had similar priorities for risk vs. baseline scenarios. Although our results are meant to illustrate the importance of considering risk, rather than directly informing real-world decisions, such shared priority areas across planning scenarios do appear to represent good return on conservation investments.

A novel contribution of our framework is that it explicitly incorporates multiple risk factors at the same time. Previous work has incorporated single risk factors analogous to those we used, including governance (Mascia & Pailler 2011; Eklund & Cabeza-Jaimejuan 2017), climate change (Hoffmann et al. 2019) and land use change (Pouzols et al. 2014; Di Minin et al. 2016) demonstrating the importance of each type of risk in protected area planning. Our results similarly demonstrate that protected area expansion decisions can be profoundly influenced by all three risk factors combined, yet they also show that relatively little additional protected area is required to account for these risks.

Despite relatively modest differences across scenarios in the amount of land required to conserve terrestrial vertebrates, shifts in the locations of priority areas sometimes resulted in substantial increases within any given nation. These asymmetries highlight the importance of cross-jurisdictional coordination to promote collaboration and improve the effectiveness of protected area systems (Dallimer & Strange 2015). In regions where nations vary widely in exposure to risks, coordination may provide opportunity to offset or otherwise mitigate risks by adjusting the geographic locations and/or boundaries of protected areas. Cooperative governance frameworks (Miller et al. 2019) are especially important for countries supporting wide-ranging species that are expected to be impacted by climate, land use, and governance risk across borders (Fig. 3). These governance frameworks would need to be developed in an environmentally just and equitable way to deliver benefits to biodiversity and local communities (Martin et al. 2013).

Within countries, the implementation of enhanced protected area networks will be highly dependent on local legal frameworks regarding land-use protection. For example, in Canada, most land (approx. 86%) is controlled by provincial and territorial governments (Neimanis 2013), although there is increasing momentum for conservation partnerships with Indigenous peoples and their governments (IUCN 2018). In contrast, in the USA the federal government is the largest landowner (controlling approx. 27% of land), with only 9% of land being owned by states (Rasker 2019 p. 20). Despite differences in governance structure among countries, cooperation among various jurisdictions within them will be essential for achieving broader protected area targets. Some level of rebalancing opportunity costs may also be necessary for those jurisdictions shouldering the largest burden of protected area expansion.

Our flexible framework and methods can also allow conservation agencies to set their own priorities from local to global scales and incorporate different metrics to assess the relevance of different forms and levels of risk. Nevertheless, we acknowledge two important caveats. First, even though we followed current practice in terms of spatial resolution for global scale analyses (Hanson et al. 2020; Jung et al. 2021) and used AOH as has been shown to be a better proxy for area of occupancy than unrefined IUCN features (Brooks et al. 2019). There is however a documented risk that processing IUCN range maps at fine spatial resolutions may overestimate biodiversity because a species is assumed to occupy all areas of a pixel (Hurlbert & Jetz 2007). Thus, our results should be assumed as maximum biodiversity estimates. Second, our estimate of the additional protected area needed to account for risk reflects the measures used in this analysis and could differ in both amount and location with other measures of risk for governance, land use or climate or if other types of risk are considered. Indeed, our risk metrics were chosen as reasonable examples, rather than definitive recommendations. Our alternative climate risk frameworks (presented in the Supporting Information) illustrate the importance of metric choice. The difference between our climate risk scenarios highlights the need for agencies to carefully consider their choices of risk metrics and suggests that smaller-scale planning exercises should choose metrics that are most relevant for each region.

The conservation community has traditionally neglected to estimate how future changes in climate (Kelly et al. 2020), land use (Di Minin et al. 2016), and governance risk might compromise the effectiveness of protected areas. Yet, as we work towards an ambitious new plan to curb biodiversity loss (CBD 2020) in a rapidly changing world, we show that incorporating future risk has profound implications for the spatial distribution of protected areas. The risk of weak governance was particularly influential. Surprisingly, incorporating risk into decision-making adds <2% to the total global area required to meet biodiversity targets. Thus, accounting for risk comes at limited extra cost which is likely outweighed by increased likelihood of achieving global biodiversity targets. Our results also emphasize the importance of cross-jurisdictional conservation initiatives, especially in adjacent countries sharing wide-ranging species where risk varies considerably from country to country. Considering risk in conservation decision-making will result in more resilient and effective conservation plans into the future to help safeguard our planet’s biodiversity in the face of the current extinction crisis.

## Data and materials availability

All data, scripts and full results are available on Open Science Framework (OSF) and will be assigned a DOI once the manuscript is in print: https://osf.io/e2fuw/?view_only=46eb2e525daf42d29df318a92762d885

## Acknowledgments

We thank Alison Johnston, Peter Arcese, Steven Cooke and Lenore Fahrig for helpful discussions. RS was funded by the Liber Ero Fellowship Program. JRB was funded by the Natural Science and Engineering Research Council of Canada (NSERC) Discovery Grant 2016-06147 and Environment and Climate Change Canada Grant GCXE19S058.

