## Supplementary Material for "Protected area planning to conserve biodiversity in an uncertain future"

**Supplementary methods:**

***Alternative climate risk measure: exposure to extreme events***

Anthropogenic climate change is affecting the frequency and duration of extreme heat events (Diffenbaugh et al. 2017; AghaKouchak et al. 2020). Exposure to these events can adversely affect human populations (Battisti & Naylor 2009; Anderson G. Brooke & Bell Michelle L. 2011; Mitchell et al. 2016) and natural systems (Harris et al. 2018; Maxwell et al. 2019). For species in natural systems, these events can further the decline and extirpation of populations, increasing the chances of extinction (Maron et al. 2015; Maxwell et al. 2019). Extreme heat events and extreme cold events can also promote the formation of novel ecosystems (Harris et al. 2018), generate enhanced selection pressures (Gutschick & BassiriRad 2003; Grant et al. 2017), and change the phenology of life history events (Sorte et al. 2016; Cremonese et al. 2017). There are a number of climate indices that have been used to estimate the occurrence of these events (Smith et al. 2013; Fenner et al. 2019). These indices are often context specific and there is little consensus on the most appropriate technique (McPhillips et al. 2018).

For this alternative measure, we estimated climatic risk based on the estimated trend in the annual proportion of days containing extreme heat events from 1979 to 2019 (La Sorte et al. 2021). Extreme heat events were estimated using hourly air temperature at 2 m above the surface and gridded at a 31 km (0.28125° at the equator) spatial resolution (Hersbach et al. 2018). The temperature data was acquired from the European Centre for Medium-Range Weather Forecasts (ECMWF) fifth generation atmospheric reanalysis of the global climate (ERA5) (Hersbach et al. 2019; Hoffmann et al. 2019 p. 5). The approach first extracted daily minimum and maximum temperature for each grid cell over the 41-year period. To reduce the influence of warming trends, the daily minimum and maximum temperature was then detrended across years for each day and grid cell using empirical mode decomposition (EMD) (Huang et al. 1998; Wu et al. 2007). The occurrence of extreme heat events was estimated using the following approach: The detrended minimum and maximum temperature data was treated as normally distributed across years for each day and grid cell. The probability density function for the detrended minimum and maximum temperature was then estimated using the mean and standard deviation calculated across years for each day and grid cell. Extreme heat events occurred when the probabilities for both minimum and maximum temperature on a given day and grid cell were within the 0.95-1.00 quartile of the probability density function. The trend in the annual proportion of days containing extreme heat events for each year was calculated for each grid cell using beta regression with a logit link function and an identity function in the precision model (Ferrari & Cribari-Neto 2004; Simas et al. 2010) (Supplementary Figures 7 – 9). See (La Sorte et al. 2021) for additional details.

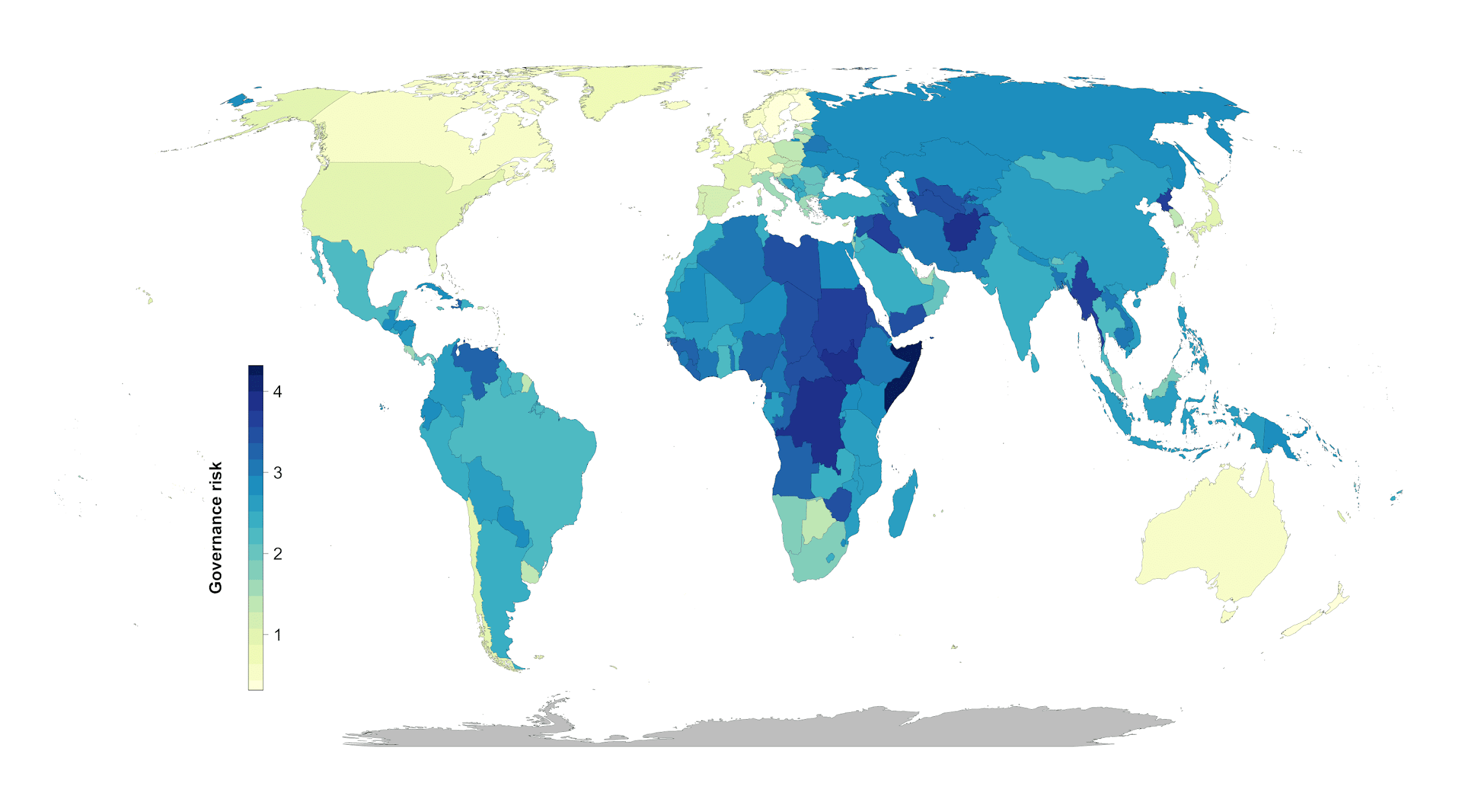

**Figure S1. Governance risk (yellow = low, blue= high)**

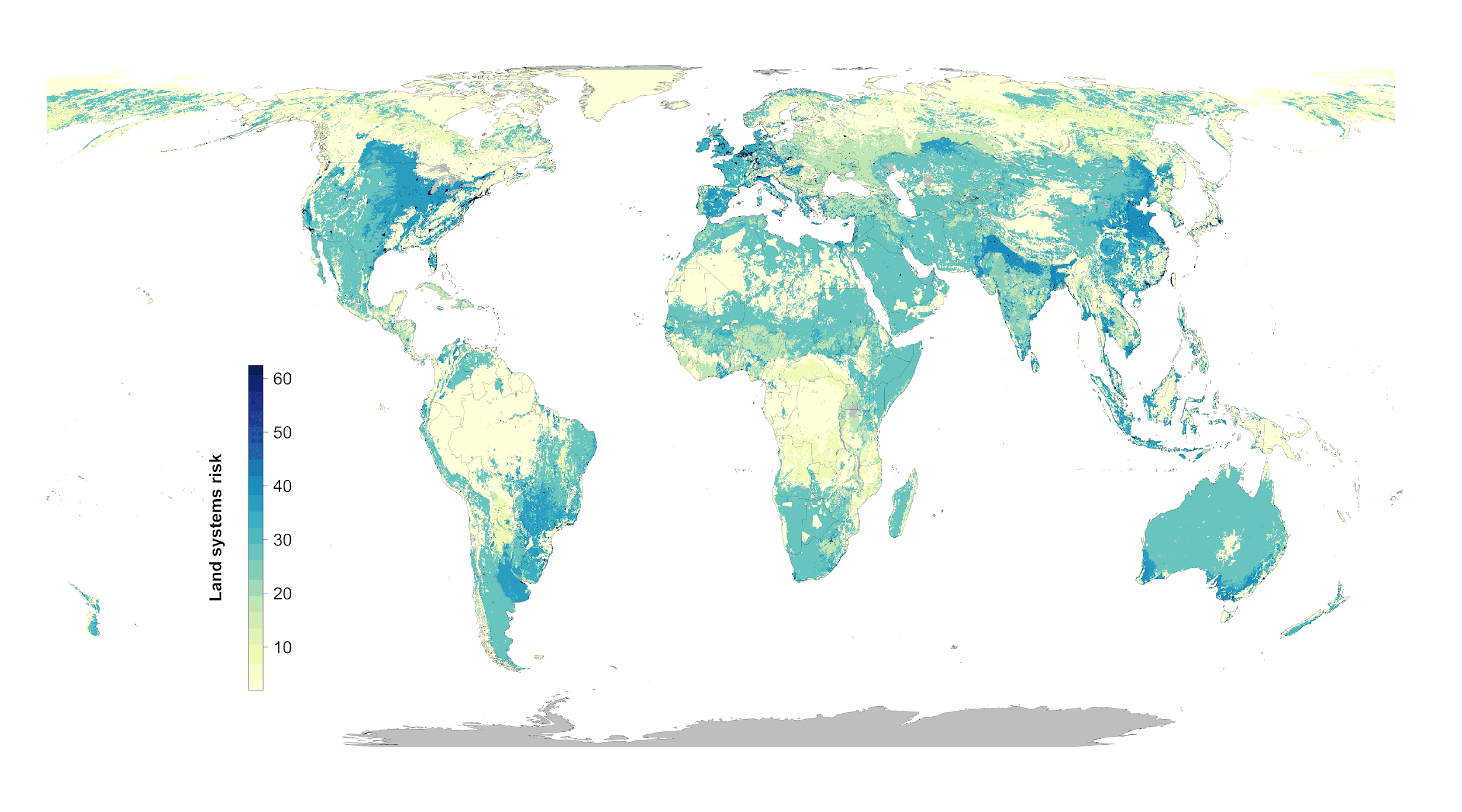

**Figure S2. Land systems risk (yellow = low, blue= high)**

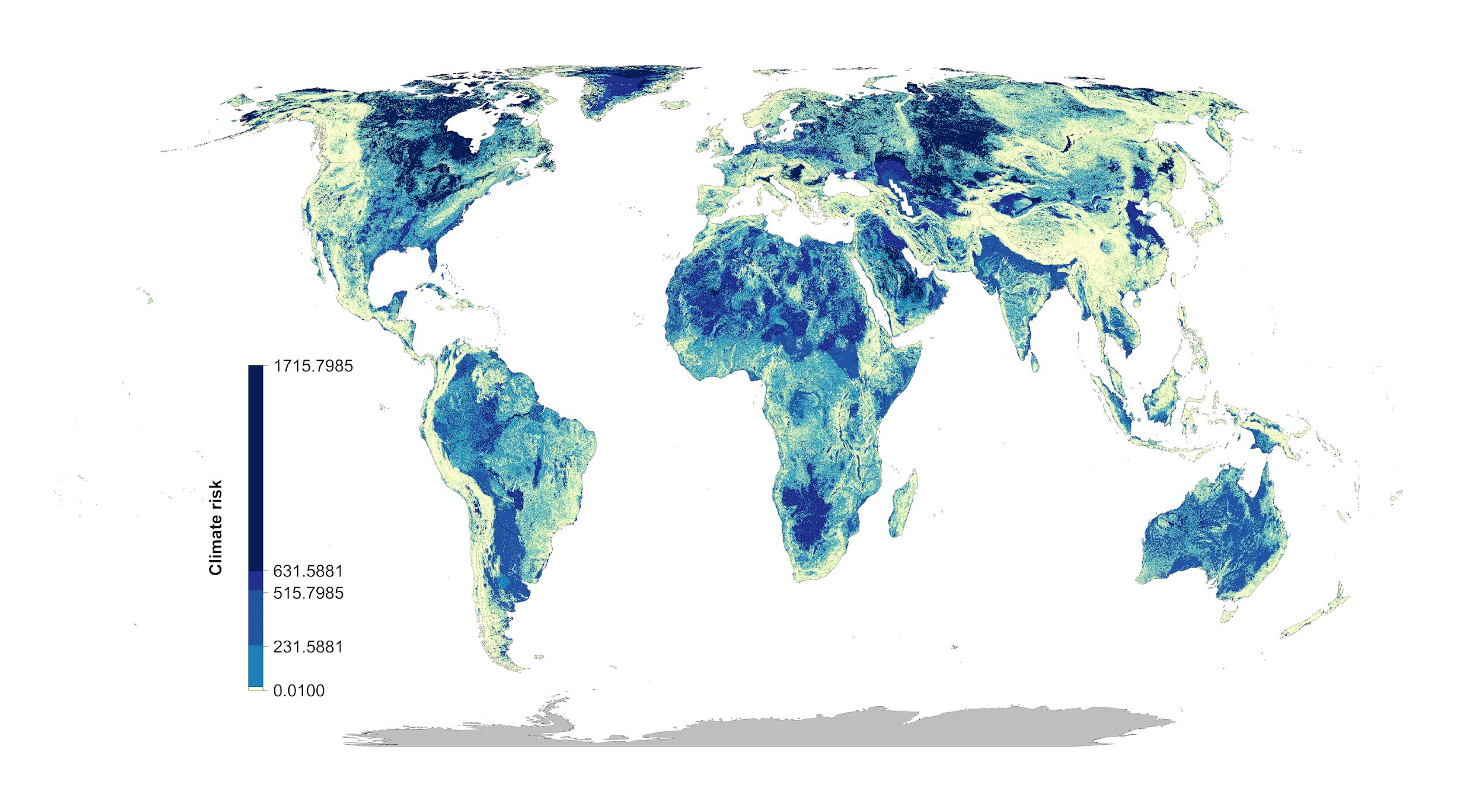

**Figure S3. Climate risk (climate velocity) (yellow = low, blue= high)**

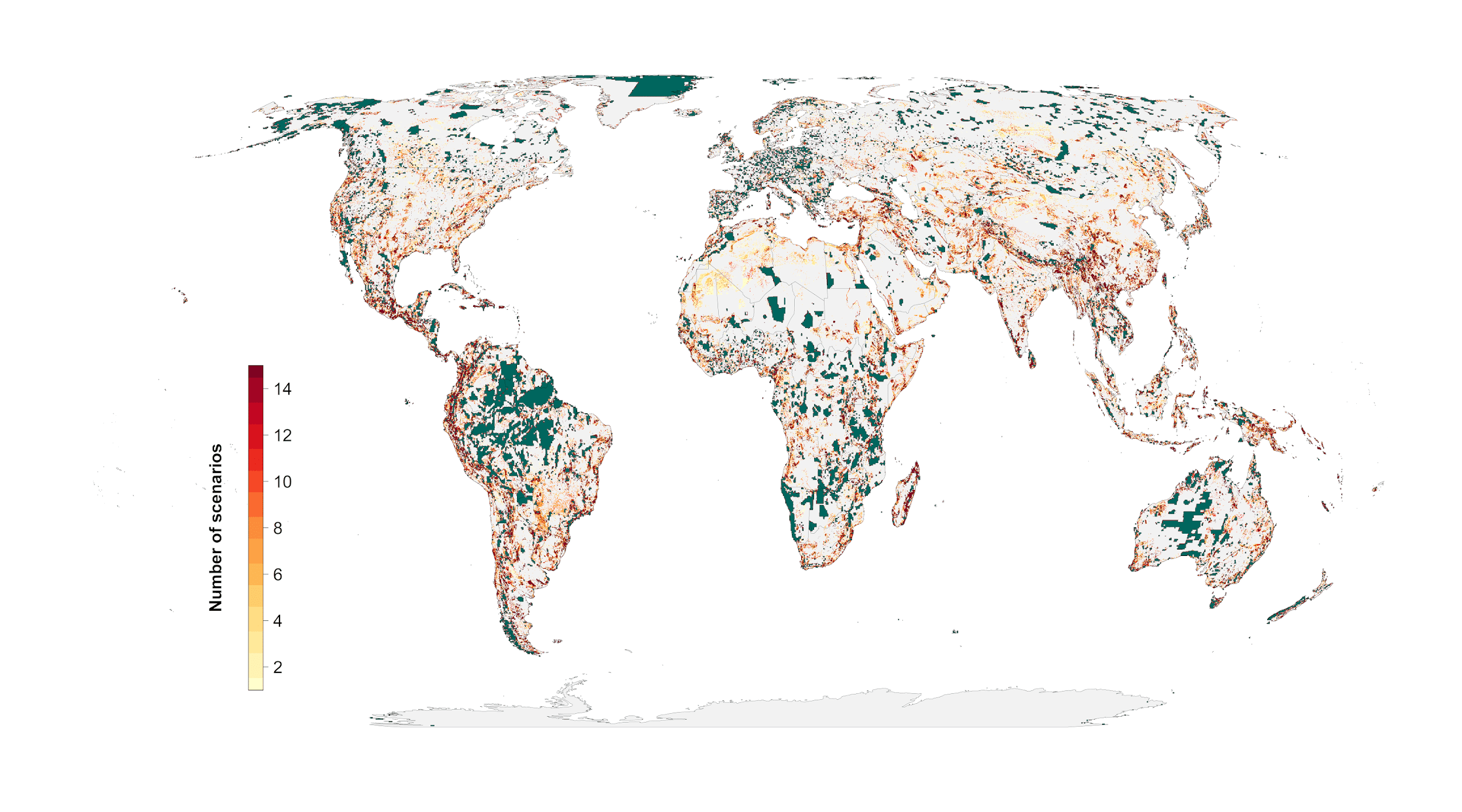

**Figure S4. Scenario overlap. green = protected areas. Color gradient from yellow (one scenario) to red (15 scenarios) = overlap.**

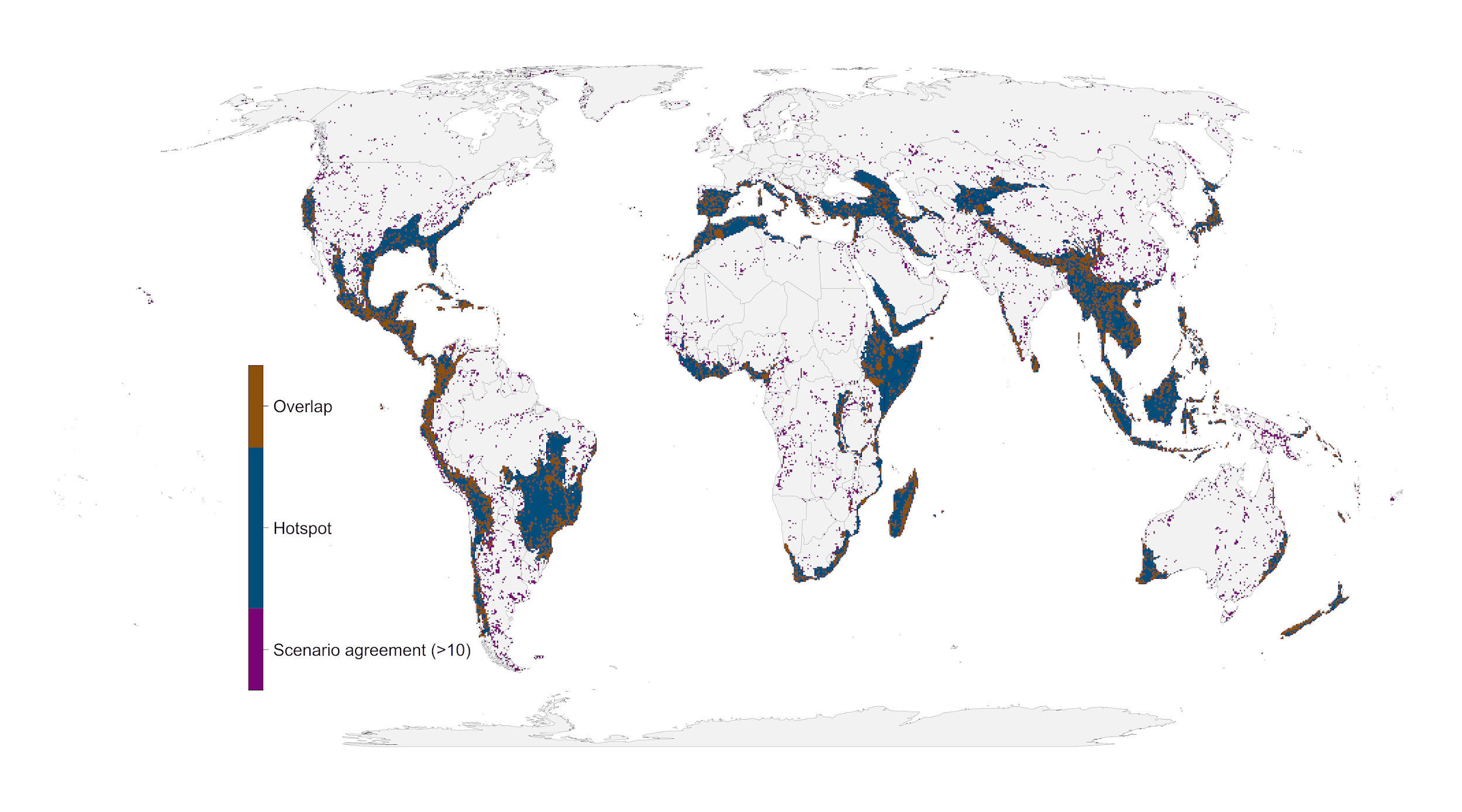

**Figure S5. Areas of high scenario overlap (>10 scenarios, green) compared to biodiversity hotspots (*28*) (blue).**

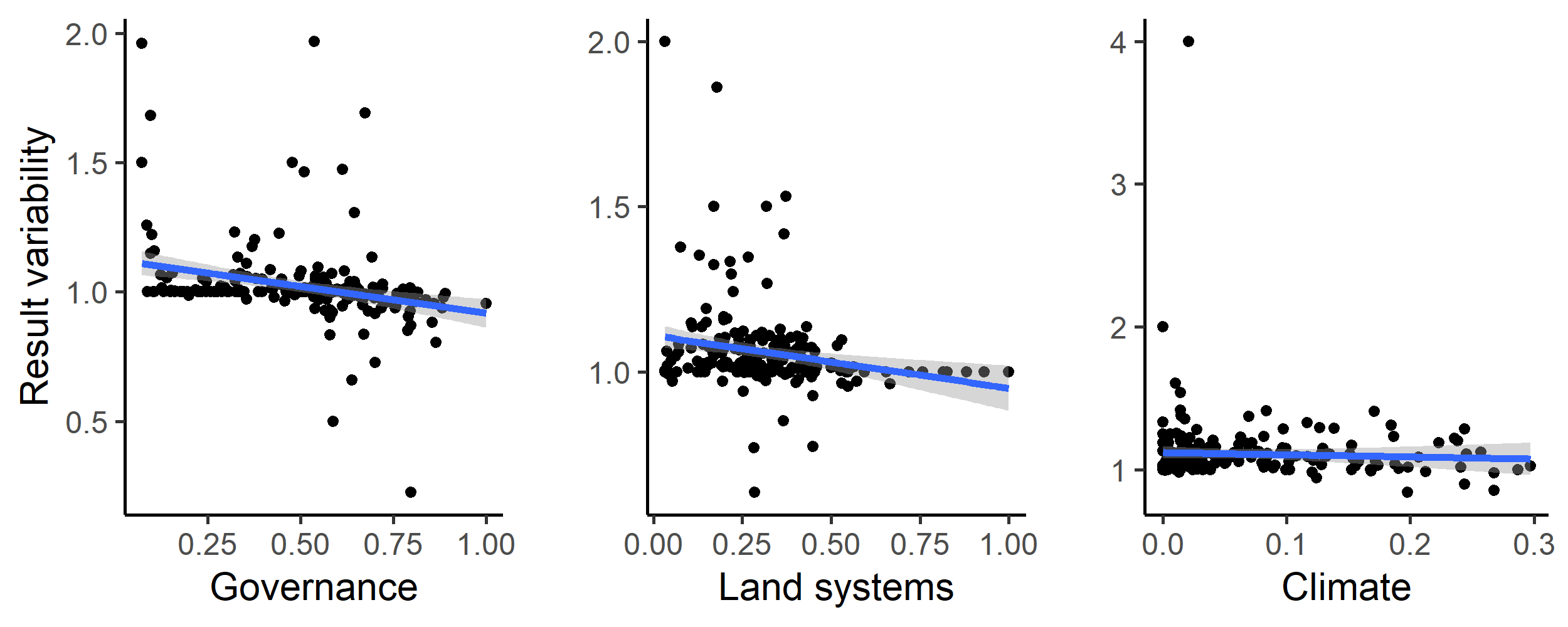

**Figure S6. Influence of average country specific risk factors on the optimization outcomes compared between null scenario and the scenarios including one of the risk factors. Each data point represents the results for one country. The fitted blue lines and 95% confidence bands are from ordinary least-squares regression.**

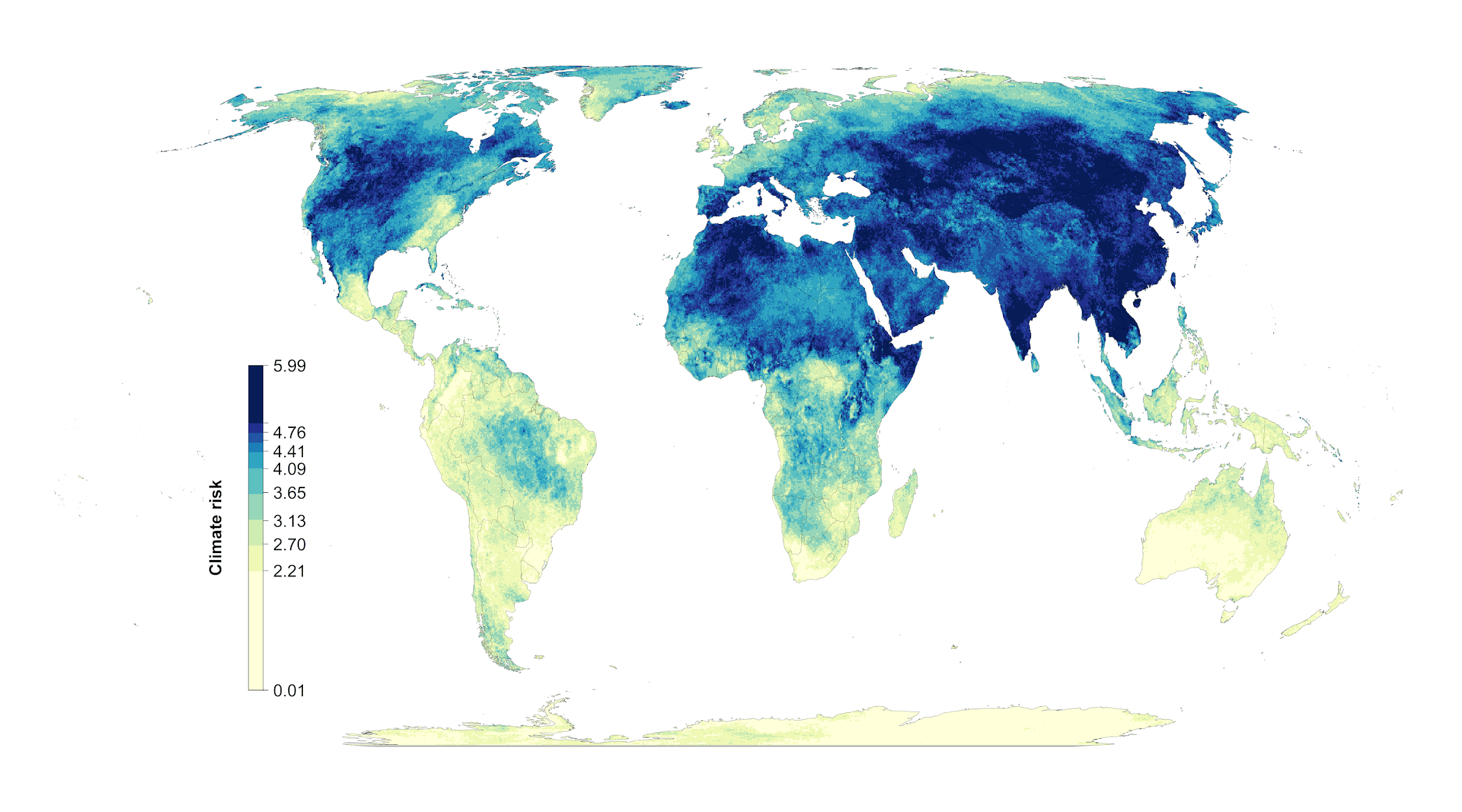

**Figure S7. Alternative climate risk metric (extreme heat events) (yellow = low, blue= high)**

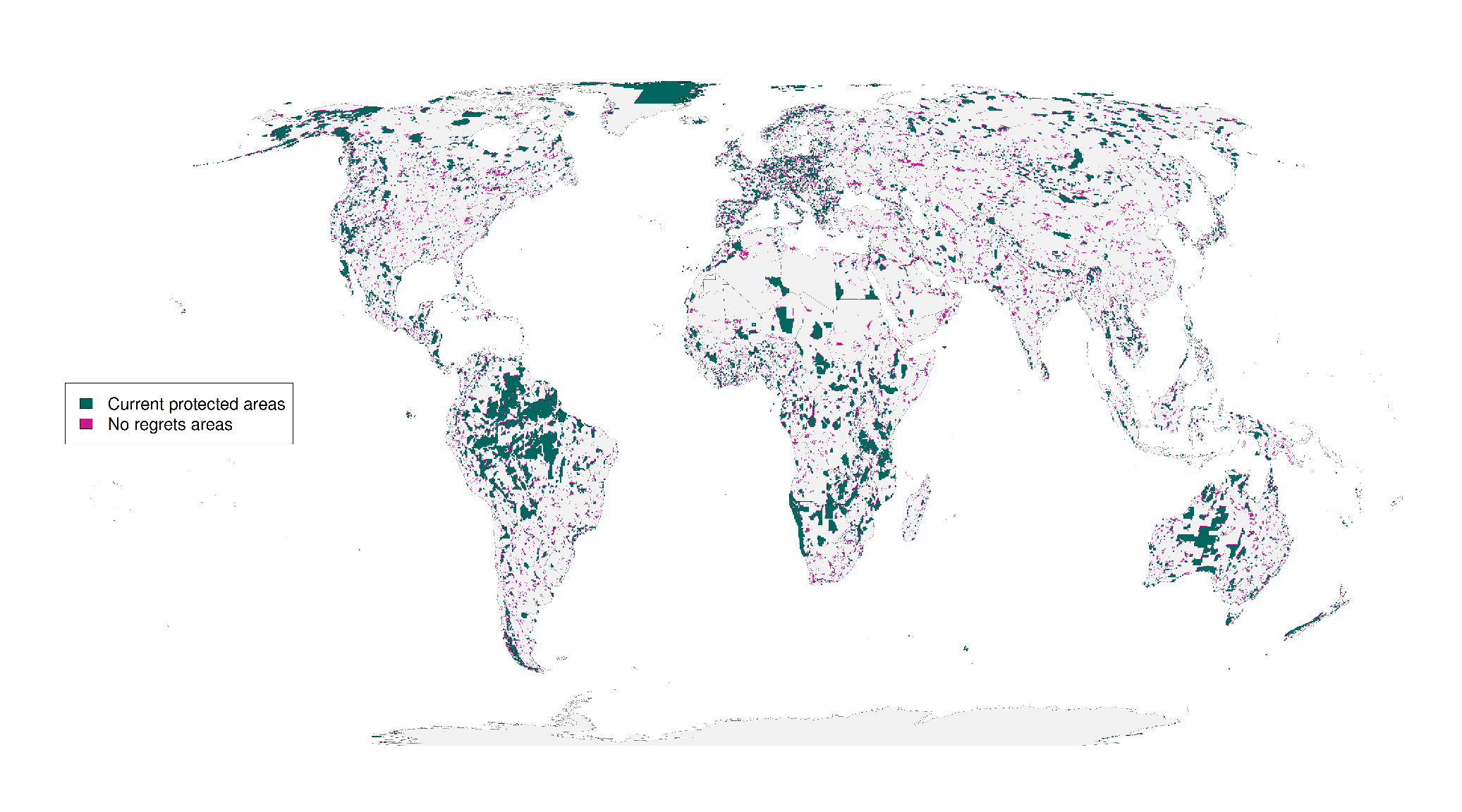

**Figure S8 Alternative climate risk scenario “No regrets” areas that were identified as priority habitat for protection regardless of the risks included in our analysis.**

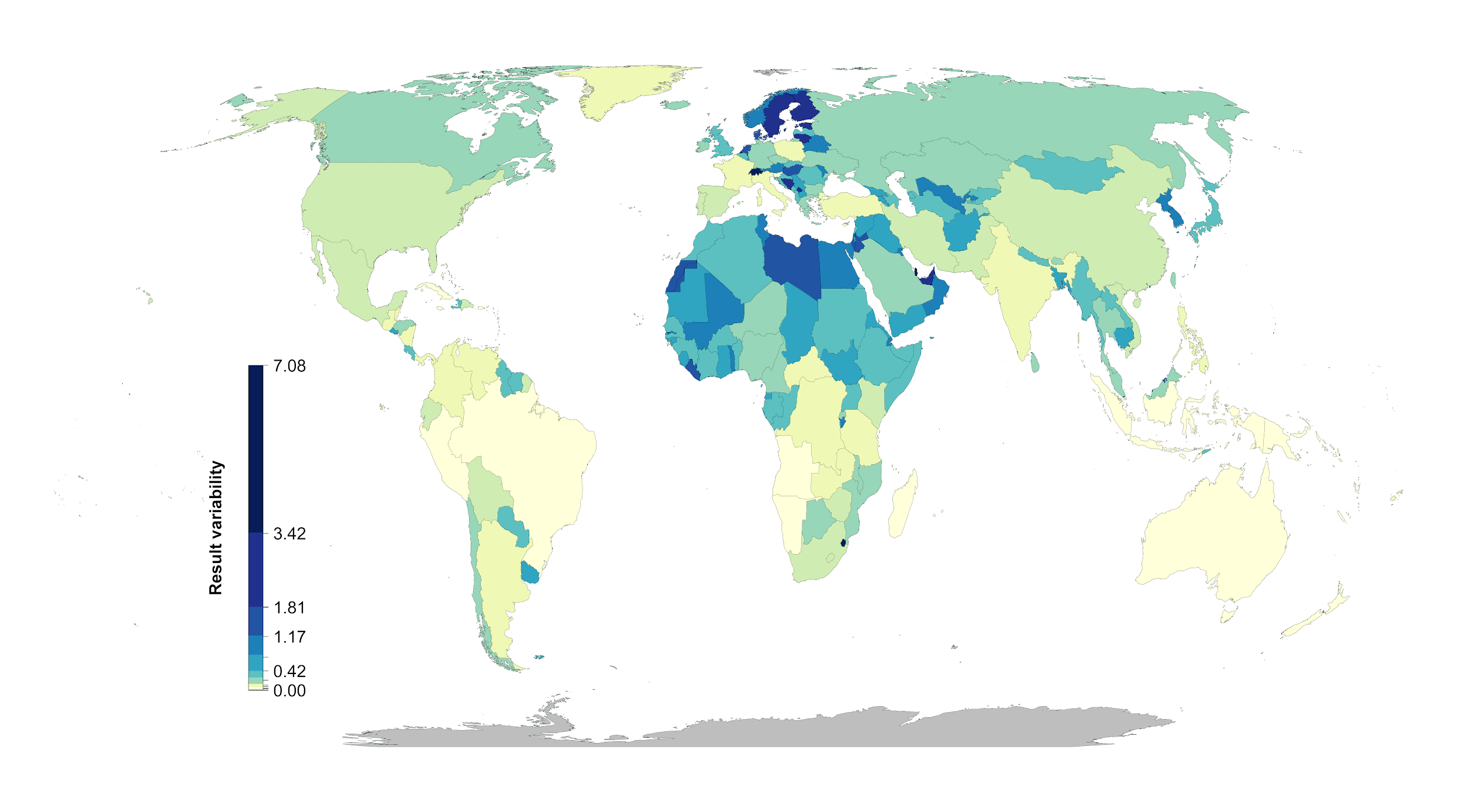

**Figure S9. Alternative climate risk scenarios percent country-level variation between the null scenario and the 15 scenarios including risk. Countries whose results are consistent across the 15 scenarios (e.g., Brazil) have low variation, while countries whose results are less consistent across the 15 scenarios (e.g., Sweden) have high variation. The kmeans method (*37*) was used to generate class intervals for visualization.**

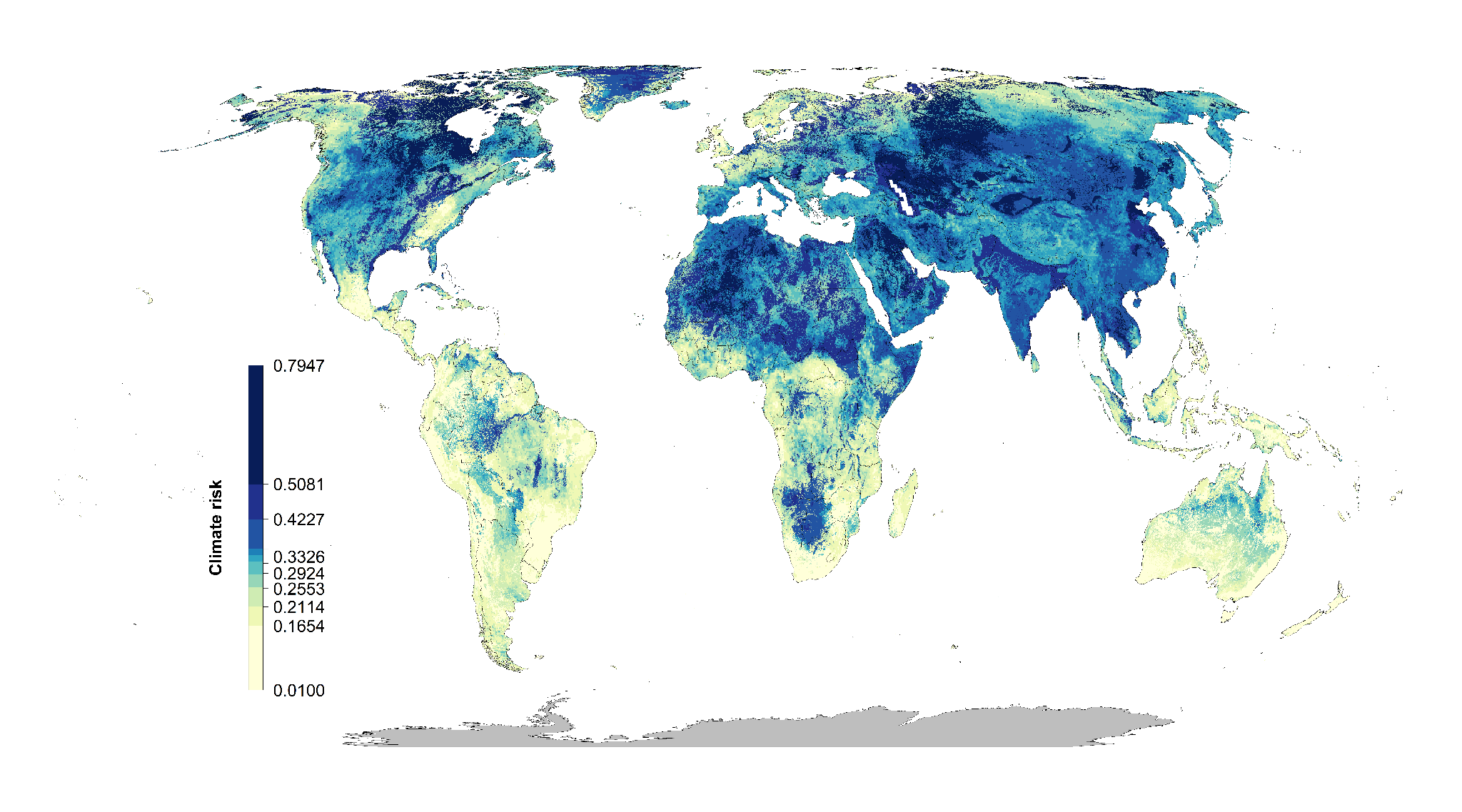

**Figure S10. Combined climate risk metric (climate velocity and extreme heat events combined) (yellow = low, blue= high)**

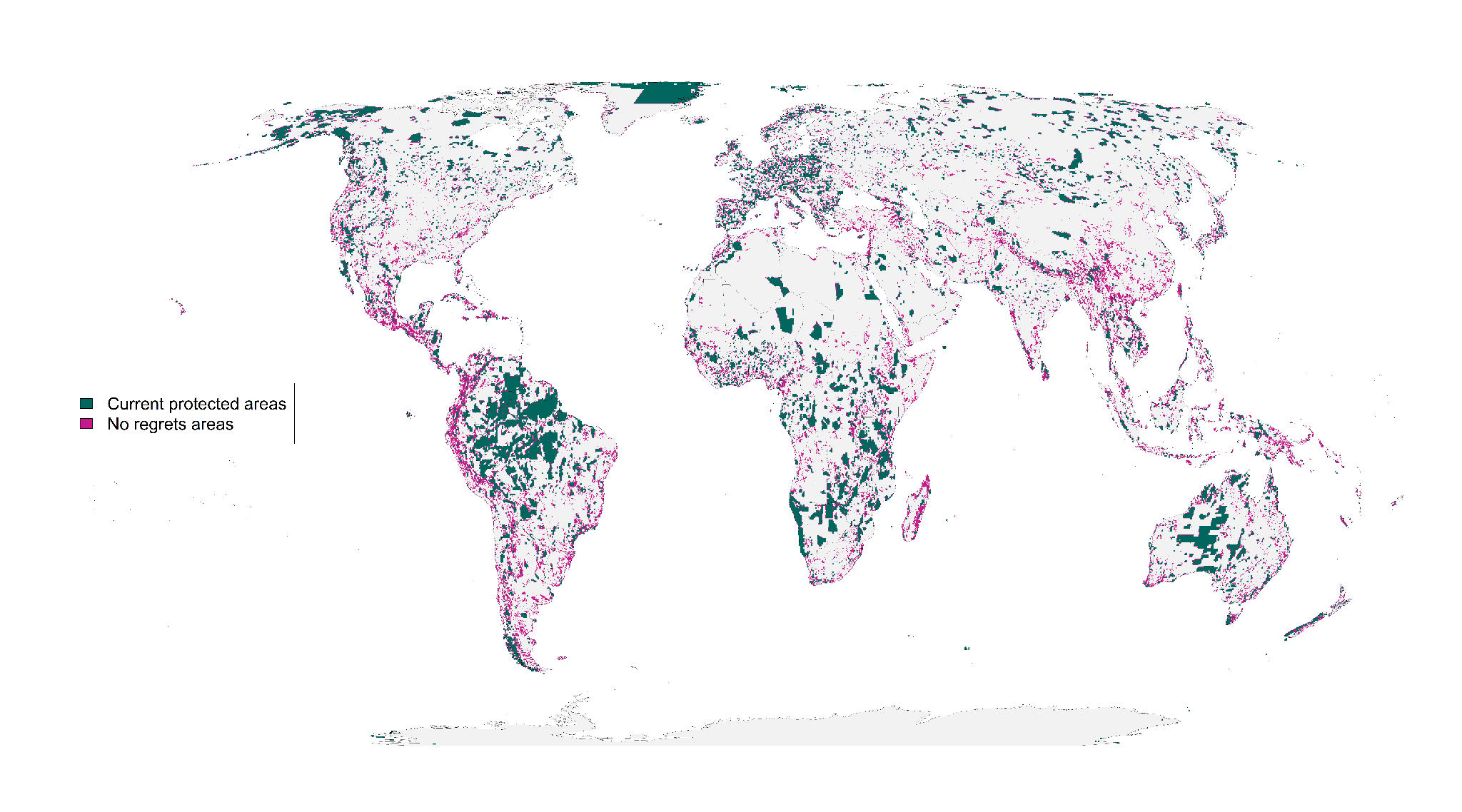

**Figure S11. Combined climate risk scenario “No regrets” areas that were identified as priority habitat for protection regardless of the risks included in our analysis.**

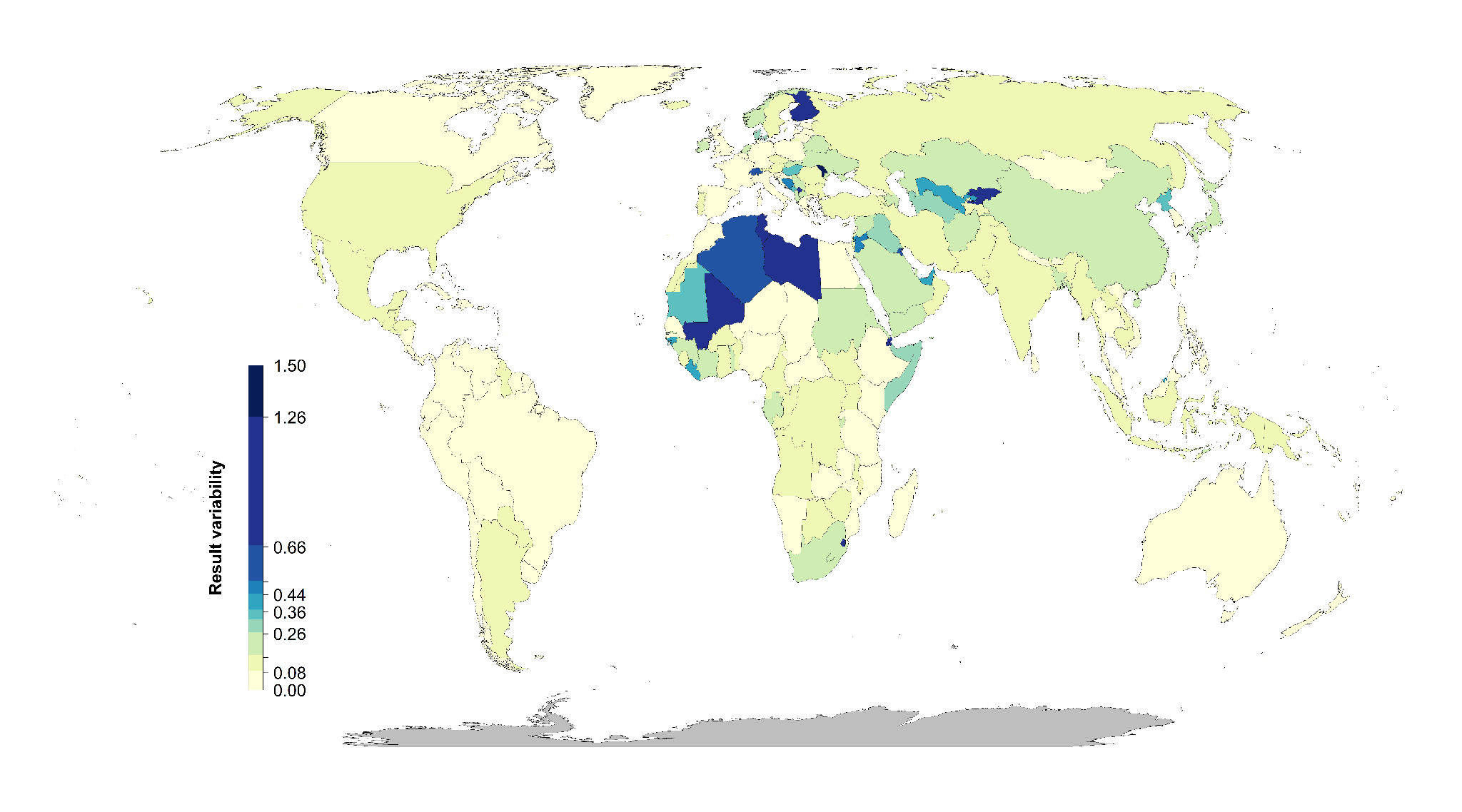

**Figure S12. Combined climate risk scenarios percent country-level variation between the null scenario and the 15 scenarios including risk. Countries whose results are consistent across the 15 scenarios (e.g., Brazil) have low variation, while countries whose results are less consistent across the 15 scenarios (e.g., Finland) have high variation. The kmeans method (*37*) was used to generate class intervals for visualization.**

**Table S1**. **Worldwide governance indicator definitions from the World Bank (15).**

| **Indicator** | **Definition**  Source: World Bank, 2020 (<https://datacatalog.worldbank.org/dataset/worldwide-governance-indicators>) |
| --- | --- |
| Voice and accountability | “Voice and accountability captures perceptions of the extent to which a country's citizens are able to participate in selecting their government, as well as freedom of expression, freedom of association, and a free media.” |
| Political stability and absence of violence | “Political Stability and Absence of Violence/Terrorism measures perceptions of the likelihood of political instability and/or politically-motivated violence, including terrorism.” |
| Government effectiveness | “Government effectiveness captures perceptions of the quality of public services, the quality of the civil service and the degree of its independence from political pressures, the quality of policy formulation and implementation, and the credibility of the government's commitment to such policies.” |
| Regulatory quality | “Regulatory quality captures perceptions of the ability of the government to formulate and implement sound policies and regulations that permit and promote private sector development.” |
| Rule of law | “Rule of law captures perceptions of the extent to which agents have confidence in and abide by the rules of society, and in particular the quality of contract enforcement, property rights, the police, and the courts, as well as the likelihood of crime and violence.” |
| Control of corruption | “Control of corruption captures perceptions of the extent to which public power is exercised for private gain, including both petty and grand forms of corruption, as well as "capture" of the state by elites and private interests.” |

**Table S2**. **Country specific results for the 15 scenarios investigated. Numbers represent % of land area of a country selected (including existing protected areas).
(As an example 5 countries included here, full list in csv. N = null, G = governance, L = land use, C = Climate)**
<https://drive.google.com/file/d/1eD4y4K8XG4nxnRL5fNtiTqzuqfIJ_DfB/view?usp=sharing>

|  | Afghanistan | Åland | Albania | Algeria |
| --- | --- | --- | --- | --- |
| N | 15.95 | 57.14 | 38.46 | 10.62 |
| G | 14.95 | 85.71 | 35.66 | 7.71 |
| L | 17.03 | 85.71 | 43.71 | 10.32 |
| C | 19.25 | 57.14 | 46.15 | 13.69 |
| GL | 15.87 | 85.71 | 37.41 | 8.94 |
| LG | 16.55 | 100 | 38.11 | 11.59 |
| GC | 19.3 | 57.14 | 46.5 | 13.71 |
| CG | 17.89 | 71.43 | 39.16 | 12.74 |
| LC | 17.8 | 71.43 | 44.06 | 13.07 |
| CL | 19.52 | 57.14 | 40.56 | 13.36 |
| GLC | 17.8 | 57.14 | 43.71 | 13.15 |
| GCL | 19.44 | 57.14 | 41.96 | 13.38 |
| LGC | 17.81 | 57.14 | 44.06 | 13.05 |
| LCG | 16.58 | 85.71 | 38.11 | 12.36 |
| CGL | 17.52 | 85.71 | 43.36 | 12.4 |
| CLG | 19.52 | 57.14 | 40.56 | 13.36 |

**Table S3. Governance risk score table (see csv)**

**(As an example Afghanistan – Barbados are included below)**
<https://drive.google.com/file/d/1g_LePBfCbphXzTiCOXCzQtNLSSYoV6me/view?usp=sharing>

| Country.Name | Country.Code | MeanIndex | SDIndex |
| --- | --- | --- | --- |
| Afghanistan | AFG | -1.65038 | 0.16074 |
| Albania | ALB | -0.28043 | 0.219515 |
| Algeria | DZA | -0.86838 | 0.121774 |
| American Samoa | ASM | 0.747997 | 0.127264 |
| Andorra | AND | 1.359029 | 0.04054 |
| Angola | AGO | -1.16429 | 0.217384 |
| Anguilla | AIA | 1.138708 | 0.225908 |
| Antigua and Barbuda | ATG | 0.687351 | 0.143042 |
| Argentina | ARG | -0.19472 | 0.196541 |
| Armenia | ARM | -0.29545 | 0.091655 |
| Aruba | ABW | 1.181311 | 0.090913 |
| Australia | AUS | 1.591282 | 0.033469 |
| Austria | AUT | 1.559385 | 0.080972 |
| Azerbaijan | AZE | -0.84662 | 0.123512 |
| Bahamas, The | BHS | 0.991142 | 0.212122 |
| Bahrain | BHR | 0.067606 | 0.189151 |
| Bangladesh | BGD | -0.8678 | 0.131258 |
| Barbados | BRB | 1.154432 | 0.145899 |
